# Semi-automatic model revision of Boolean regulatory networks: confronting time-series observations with (a)synchronous dynamics

**DOI:** 10.1101/2020.05.10.086900

**Authors:** Filipe Gouveia, Inês Lynce, Pedro T. Monteiro

## Abstract

**Motivation:** Complex cellular processes can be represented by biological regulatory networks. Computational models of such networks have successfully allowed the reprodution of known behaviour and to have a better understanding of the associated cellular processes. However, the construction of these models is still mainly a manual task, and therefore prone to error. Additionally, as new data is acquired, existing models must be revised. Here, we propose a model revision approach of Boolean logical models capable of repairing inconsistent models confronted with time-series observations. Moreover, we account for both synchronous and asynchronous dynamics.

**Results:** The proposed tool is tested on five well known biological models. Different time-series observations are generated, consistent with these models. Then, the models are corrupted with different random changes. The proposed tool is able to repair the majority of the corrupted models, considering the generated time-series observations. Moreover, all the optimal solutions to repair the models are produced.

**Contact:** {}

## 1 Introduction

Complex cellular processes can be represented through biological regulatory networks which describe interactions between biological compounds such as genes and proteins. Having a computational model of such networks is of great interest, allowing the reproduction of known behaviours, the test of hypotheses, and the identification of predictions *in silico*.

Different formalisms have been used by the community to model biological regulatory networks, such as Ordinary Differential Equations (ODE), Piecewise-Linear Differential Equations (PLDE), Logical Formalism, Sign Consistency Model (SCM), among others (De Jong, 2002; Karlebach and Shamir, 2008; Siegel *et al.*, 2006). Here, we consider the (Boolean) logical formalism, which has proven useful to study and analyse dynamical behaviour of biological models (Thomas, 1973; Naldi *et al.*, 2015; Abou-Jaoudé *et al.*, 2016; Selvaggio *et al.*, 2020).

As these models are extended or new experimental data is acquired, it becomes necessary to assess whether the models continue to be consistent with the existing data, and if necessary, to revise them. Typically, model construction and model revision is a manual task, performed by a domain expert, and therefore prone to error. Moreover, there is an inherent combinatorial problem associated to all possible changes to render a model consistent. This calls for automated approaches to build and revise such models, especially when incomplete information is considered. The notion of *belief revision* exists for a long time and addresses the problem of having new information in conflict with previously known information (Alchourrón *et al.*, 1985). Even though model revision has been addressed by the community for models of regulatory networks (Gebser *et al.*, 2010; Guerra and Lynce, 2012; Merhej *et al.*, 2017), few approaches have been proposed for (Boolean) logical models (Mobilia *et al.*, 2015; Lemos *et al.*, 2019).

In this work, we propose a model revision approach for Boolean logical models considering time-series observations. Previous works only considered repairing inconsistent Boolean logical models under stable state observations (Gouveia *et al.*, 2019a,b). Here, we tackle a complementary problem taking into account the model dynamic behaviour, under both synchronous and asynchronous update schemes.

The paper is organised as follows. Section 2 introduces key concepts relevant for this work. An overview of the related work is given in Section 3. The proposed approach is presented in detail in Section 4. Section 5 details the experimental evaluation and shows the obtained results. This paper concludes with Section 6 and future prospects are presented.

## 2 Preliminaries

Biological regulatory networks describe complex biological processes. A regulatory network is composed of a set of biological compounds, like genes and proteins, and the set of interactions between them. Modelling such networks is particularly useful to be able to computationally reproduce existing observations, test hypotheses, and identify predictions *in silico*.

Typically, a regulatory network is represented by a directed graph *𝒢* = (*V, E*), known as *regulatory graph*, where *V* is the set of nodes, representing the set of compounds, and *E ⊆ {*(*u, v*): *u, v ∈ V}* is the set of directed edges, representing the set of regulatory interactions. If (*u, v*) *∈ E*, then we say that *u* is a *regulator* of *v*. It is common to associate a sign to each edge, representing a positive interaction (activation) or a negative interaction (inhibition).

Although these regulatory graphs define the structure or *topology* of the network, they still need information about the regulatory rules to properly define a computational model, capable of describing the system’s dynamics. Different formalisms have been used to describe such models (see Karlebach and Shamir (2008) for a detailed review). In this work, we focus on the Boolean logical formalism presented by Thomas (1973).

### 2.1 Logical formalism

A logical model of a regulatory network is defined by a tuple (*V, 𝒦*), where *V* = *{v*_1_, *v*_2_, *…, v*_*n*_} is the set of *n* variables representing the regulatory compounds of the network, where to each *v*_*i*_ is assigned a non-negative integer value in {0, *…, max*_*i*_}, representing the compound concentration level. In this work, we consider Boolean logical models, where 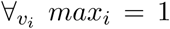, *i.e.*, each compound of the network is represented by a Boolean value, meaning that the compound is either present (active) or absent (inactive). The set of *n* regulatory functions *𝒦* = *{K*_1_, *K*_2_, *…, K*_*n*_} are, in this case, a set of Boolean functions over *V*. Let ℬ be the set {0, 1} and ℬ^*n*^ be the *n*-dimensional Cartesian product of the set ℬ. A Boolean function *f* is then defined as *f*: ℬ^*n*^ *→ ℬ*.

A Boolean function can be represented in Disjunctive Normal Form (DNF) as a disjunction (OR) of terms, where each term is a conjunction (AND) of literals and each literal is a variable or its negation (Biere *et al.*, 2009). However, a Boolean function can have multiple equivalent DNF representations. To uniquely represent a Boolean function, the Blake Canonical Form (BCF) can be used. BCF is a special case of DNF where a Boolean function is represented by the disjunction of all its *prime implicants* (for details, see Crama and Hammer (2011)). As a Boolean function is uniquely identified by the list of its prime implicants, any given Boolean function has a unique BCF representation. In this work, we restrict the domain of the Boolean regulatory functions to the set of monotone non-degenerate Boolean functions.

Given a monotone Boolean function in the BCF representation, each variable *x*_*i*_ always appears either as a positive (*x*_*i*_) or as a negative (*¬x*_*i*_) literal. In the context of biological regulatory networks, this means that a given regulator *v*_*i*_ of *v*_*j*_ only has one role, either always appearing as a positive variable in the regulatory function of *v*_*j*_ (an activator), or always appearing as a negated variable (an inhibitor).

A non-degenerate Boolean function is a function that depends on all of its variables, *i.e.*, all variables have an impact on the function result (Wegner, 1985). In the context of biological regulatory networks, this means that in each regulatory function only its regulators are present as variables of the function. If a given variable does not affect in any way the output of the regulatory function, then the variable should not be considered a regulator.

Figure 1 illustrates a Boolean logical model where green pointed arrows represent positive interactions (activations), and red blunt arrows represent negative interactions (inhibitions). On the right side of the graph are represented the regulatory functions. Note that *v*_4_ does not have any regulator, and therefore is considered an *input node* (having no associated regulatory function). It is also possible to verify that each regulator appears with the correspondent sign on the regulatory functions according to the type of interaction. For example, *v*_1_ is a positive regulator of *v*_3_ and appears as a positive variable in the regulatory function of 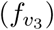. In contrast, *v*_4_ has a negative interaction with *v*_3_, given that *v*_4_ appears as a negative literal in 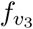.

**Figure 1:**
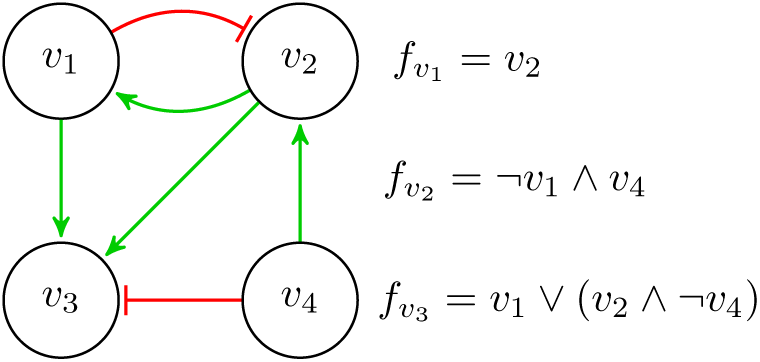
Example of a Boolean logical model.

### 2.2 Dynamics

Computational models of biological regulatory networks are particularly useful to analyse and simulate the state of the network over time. The state of a network can be represented by a vector *S* = [*v*_1_, *v*_2_, *…, v*_*n*_], where *v*_*i*_ represents the value of the variable associated with the *i*-th compound of the network. The generated dynamic of a network can be represented as a State Transition Graph (STG), where each node represents a state of the network, and directed edges between nodes represent possible transitions between states. Considering a Boolean logical model of a regulatory network with *n* nodes, where each node can only have one of two possible values, there are 2^*n*^ different states of the network. Therefore, there is an exponential state explosion with the increase of the size of the network.

The dynamics of a regulatory model can be generated according to different update schemes, such as synchronous or asynchronous updates. In the synchronous update scheme, all the regulatory functions are applied simultaneously, updating the value of all the corresponding variables. In contrast, in the asynchronous update, only one regulatory function is applied at each time, meaning that only one compound has its value updated. Different update schemes generate different transitions between states of the STG. The impact of the update scheme in the STG lies in the number of transitions, where in the synchronous update only one transition exits from each state, while in the asynchronous update several transitions may exist.

There is a combinatorial explosion of the generated dynamics when the asynchronous update scheme is considered. Since only one regulatory function is applied at each time step, multiple choices can be made, and therefore different dynamics can be generated. This makes the analysis and study of dynamics a difficult task, in particular if the asynchronous update scheme is considered. However, the asynchronous update scheme is more realistic than the synchronous update, due to the fact that biological processes do not naturally occur in a synchronous way.

An important property to be verified in a network dynamic is the existence of *attractors*. An attractor is a set of states of the network such that: from each state belonging to the attractor it is possible to reach any other state of that attractor through one or more state transitions; and there is no state transition from any state of the attractor to a state outside the attractor. These attractors correspond to the *terminal* Strongly Connected Components (SCC) of an STG. An attractor comprised by only one state is denoted a *point attractor (or stable state)*, corresponding to an equilibrium state of the network. Otherwise, it is denoted a *complex or cyclic attractor*, representing changes between the states of the network in a cycle or loop.

## 3 Related Work

Computational models of biological regulatory networks are essential to study the behaviours of the correspondent regulatory networks. Different problems arise with the study of such models, with different approaches to tackle each problem being proposed in Systems Biology, from model building to model analysis, as well as model revision.

From experimental data it is possible to infer the interactions between compounds, as well as their corresponding regulatory functions. Some of the difficulties of this task are, the lack of experimental data, leading to several model possibilities, or the existence of incorrect or noisy data that leads to incorrect biological models. Moreover, when considering Boolean logical models, it is necessary to take into account that there are at most 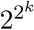 candidate Boolean functions with *k* regulators, since several functions may be consistent with the experimental data available for each compound, and a choice must be made. To infer one possible model, statistical techniques are commonly used (Butte and Kohane, 1999; Friedman, 2004). Some approaches, instead of determining the most likely model from experimental data, try to enumerate all feasible models (Guziolowski *et al.*, 2013; Ostrowski *et al.*, 2016).

The study of a model, in particular the study of its generated dynamic behaviours and the comparison between such behaviours and experimental observations, is of great interest. Due to the STG combinatorial explosion several formal verification techniques have been proposed to analyse dynamic behaviour. Some of these techniques include model-checking approaches to automatically verify reachability properties (Monteiro *et al.*, 2014; Ostrowski *et al.*, 2016). Reachability verification addresses the problem of checking whether, from a given state, it is possible to reach a given target state through valid state transitions. This is particularly important to verify if a model is able of generate and justify some observed behaviour, or to predict how a network can evolve from a specific state. Model reduction techniques to reduce the size of the generated dynamics have also been proposed (Naldi *et al.*, 2011; Pauleve, 2018). These techniques aim to reduce the combinatorial explosion of the state space when analysing dynamical properties of a network. Since attractors are a key property of interest of the network dynamics, some approaches have been proposed to determine attractors under synchronous and asynchronous update schemes (Fayruzov *et al.*, 2011; Hopfensitz *et al.*, 2012).

When new experimental data is acquired, it is necessary to validate if the computational model is consistent with the new information, *i.e.*, if the computational model can reproduce the same experimental data. However, most models are manually built by a modeller (a domain expert) and therefore are prone to error. Even considering automated approaches to built such models, only known and most likely incomplete information is considered, and an incorrect model may still be generated (although consistent with the knowledge at the time). Given new information, a model may no longer be consistent, and therefore it should be revised and updated. The notion of *belief revision* exists for a long time and addresses the problem of having new information in conflict with previously known information (Alchourrón *et al.*, 1985). However, there are few approaches to model revision of Boolean models of biological regulatory networks.

Model revision approaches were proposed over the Sign Consistency Model (SCM) formalism (Siegel *et al.*, 2006; Gebser *et al.*, 2010; Guerra and Lynce, 2012). The SCM relies on the difference between concentration levels, being different from the Boolean logical model. In the SCM, each variable has value “+” (resp. “−”) if it represents an increase (resp. decrease) in the concentration level of the biological compound. Moreover, the regulation of each compound is based on sign algebra, being the sum of the products between the value of each regulator and the sign of the corresponding interaction. This type of models lack expressiveness of the regulatory functions compared with the logical model.

A first approach to model revision over Thomas’ logical formalism (Thomas, 1973) was proposed by Mobilia *et al.* (2015). In this work, it is considered that a model is inconsistent when it is over-constrained. Therefore, to repair an inconsistent model, it is considered the removal of some constraints, under a minimisation criterion, until the model is no longer inconsistent. This may lead to under-constrained models, not correctly representing the corresponding biological process.

Model revision usually operates under a minimal assumption as there can be several ways to repair a model. Most approaches define atomic repair operations and minimise the number of operations applied to render a model consistent. Merhej *et al.* (2017) proposed the use of rules of thumbs, which are properties found in the literature, in order to repair inconsistent Boolean models. This work considers Boolean models where the regulatory function of each compound is as follows: it becomes active if there is at least one active activator and there is no active inhibitor; it becomes inactive if there is at least one active inhibitor and there is no active activator; otherwise its value is not changed. When repairing such models, the synchronous update scheme is considered, and the possible repair operations consist in adding or removing edges between nodes of the model.

More recently, Lemos *et al.* (2019) proposed a model revision procedure over Boolean regulatory networks, where more expressive regulatory functions are considered when compared with the previously mentioned approaches. Boolean functions are considered as regulatory functions, represented by the combination of three basic Boolean functions (AND, OR, and IDENTITY). Moreover, the set of defined repair operations comprise: changing the type of interaction of a regulator; changing a Boolean operator from OR to AND and vice-versa; and removing a regulator. This approach does not consider the impact that changing a regulatory function has on the network dynamics. Moreover, regulatory functions are defined in a gate-like manner, where changes operate over AND and OR operators. As a consequence, not all the possible Boolean functions are considered when repairing an inconsistent model.

## 4 Approach

In previous works, we have successfully addressed the model revision problem considering only stable state observations (Gouveia *et al.*, 2018, 2019a,b). Here, we expand some of the techniques of these works to be able to consider the model dynamics and time-series observations. The proposed approach was implemented as a tool freely available online^1^.

In this work, we propose a new model revision approach of Boolean models that considers time-series observations and the network’ (a)synchronous dynamics, addressing some of the limitations of previous approaches (Section 3). In particular, we consider monotone nondegenerate Boolean functions as regulators, allowing for more expressive regulatory functions than some of the previous approaches (Siegel *et al.*, 2006; Gebser *et al.*, 2010; Guerra and Lynce, 2012; Merhej *et al.*, 2017). We define an optimisation criteria for possible model repairs which takes into account how these models are built, and the different degrees of confidence in the correctness of model components (interactions and regulatory functions). Moreover, we define four atomic repair operations (detailed below), and take into account the impact of changing the regulatory functions on the model dynamics.

To represent a Boolean logical model, we consider the Answer Set Programming (ASP) language due to its expressive power. Moreover, ASP has been used to represent biological models and proved useful in model inference, consistency check, and model revision approaches (Siegel *et al.*, 2006; Gebser *et al.*, 2010; Merhej *et al.*, 2017).

To define the regulatory graph, we use a predicate `vertex(V)`, to indicate that V is a vertex of the graph, and a predicate `edge(V1,V2,S)` to represent an edge from `V1` to `V2` with a sign `S` *∈ {*0, 1}. Note that a vertex represents a biological compound and an edge represents an interaction, where a positive sign (1) indicates an activation, and a negative sign (0) represents an inhibition. To represent monotone non-degenerate Boolean functions in BCF, we define the predicate `functionOr(V,1..N)` that indicates that the regulatory function of V is a disjunction of N terms. Then we define the predicate `functionAnd(V,T,R)` representing that the compound R is a regulator of V and is present in the term T of the regulatory function.

Considering the predicates described above, the model represented in Figure 1 is defined as follows.

**Listing 1:**
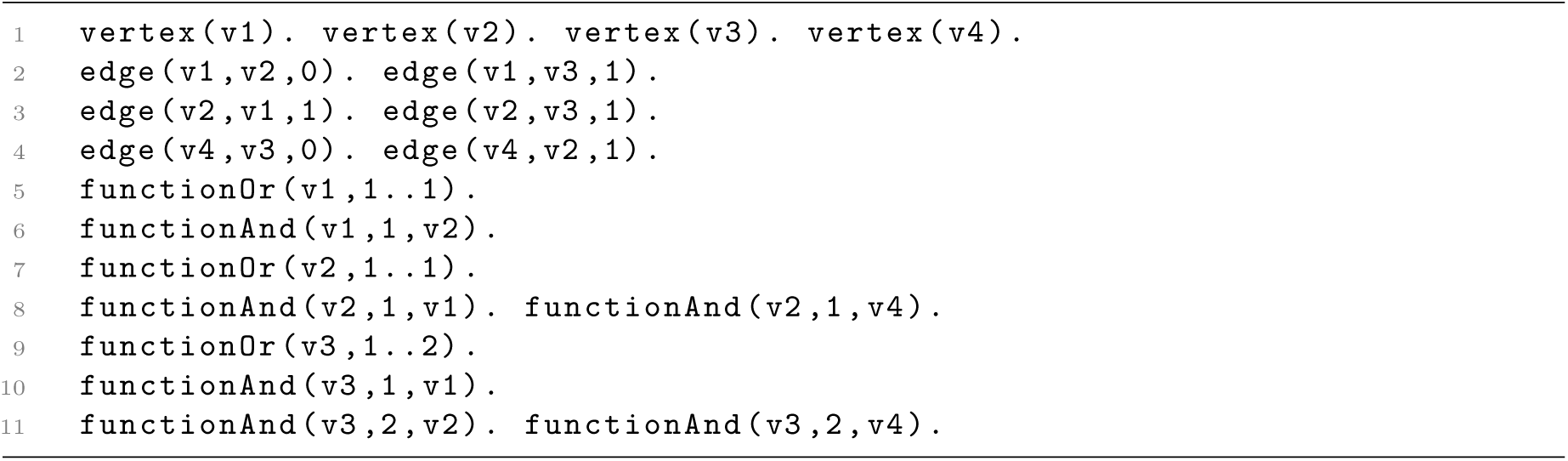
Input example for the model in Figure 1.

Note that it is not necessary to explicitly define the predicate vertex as this information can be inferred from the predicate `edge`. Also note that the sign of each literal is not represented in the predicate `functionAnd` as this information is on the edge predicate.

The experimental observations are represented as time-series data, with discrete time steps. To encode time-series observations, the predicate `exp(E)` identifies an experimental observation E. Then, to represent the observed values of an experiment, we use the predicate `obs vlabel(E,T,V,S)`, which means that in experiment E node V has an observed value S *∈ {*0, 1} at time T.

Figure 2 illustrates the proposed approach architecture. Given a model and a set of experimental observations, the goal is to repair the model if needed. To do so, first the consistency of the model is verified (detailed in Section 4.1). If the model is consistent, then no repair is needed. Otherwise, a set of repair operations must be applied. The computation of the set of repair operations that render a model consistent is described in detail in Section 4.2. It may be possible to repair an inconsistent model with different sets of repair operations. Our tool produces all possible sets of repair operations that can be applied under a defined optimisation criterion.

**Figure 2:**
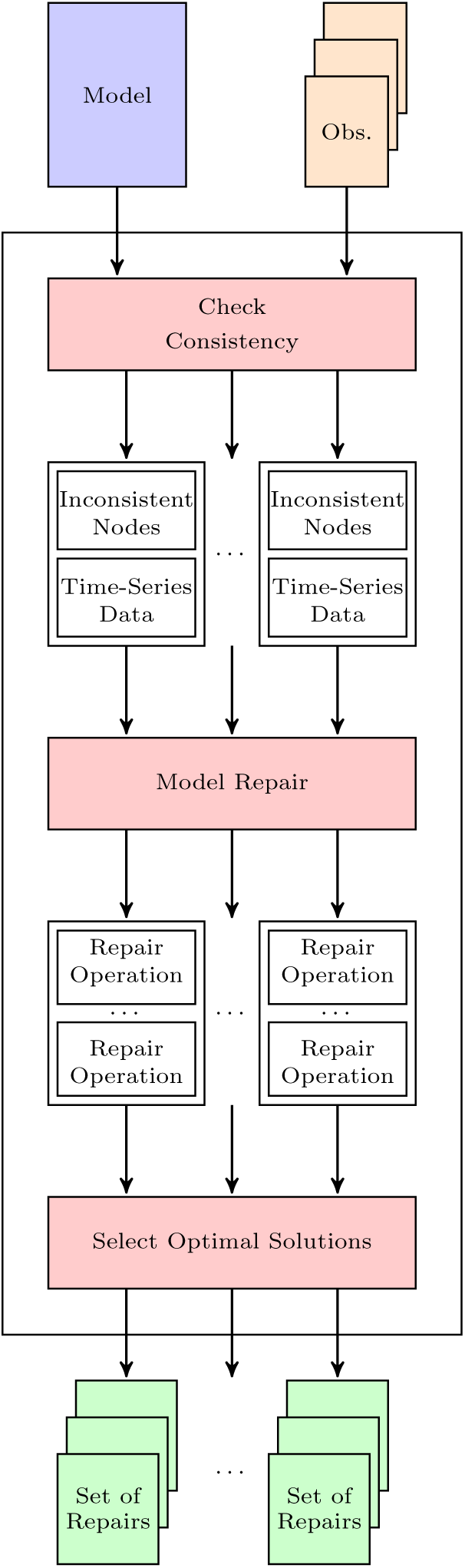
Approach architecture. The tool receives as input a model and a set of observations. It verifies the consistency, producing all the minimum possibilities of inconsistent nodes and corresponding complete time-series data. For each possible inconsistency, the minimal set of repair operations are computed (if exist). The optimal set of repairs are selected and produced as output.

### 4.1 Consistency Check

Given a model and a set of experimental observations, it is necessary to first check the consistency of the model, as shown in Figure 2. If the model is consistent with the observations, then no repair is necessary. Otherwise, if the model cannot reproduce the experimental observations, it is necessary to identify the possible reasons of inconsistency in order to repair the model. We say that a node is *inconsistent* if its regulatory function cannot reproduce, at some time step, the correspondent observed value, given the observed values of its regulators at the previous time step. Considering the synchronous update, every regulatory function is applied at each time step. Considering the asynchronous update, only one regulatory function is applied at each time step, and a node is inconsistent if the regulatory function is applied at that time step and it does not reproduce the observed value.

We use an ASP approach to verify the consistency of a model given a set of experimental observations. In order to achieve this, we define a set of rules. As we may have incomplete time-series data, *i.e.*, missing observed values in the experimental observations, a value must be assigned to each node. We use the predicate `vlabel` to assign a value to each node, with the same notation as the `obs vlabel` predicate (line 4 of Listing 2). Then, in line 5, we define that it may not exist an assigned value different from the observed value. This ensures that if we observe a value at a given time step, then it should correspond to its expected value. Moreover, this generates all the possible combinations of missing observed values, and a complete time-series data is generated. Having a complete time-series data when missing values exist is important to the repairing process, as will be described later in Section 4.2.

**Listing 2:**
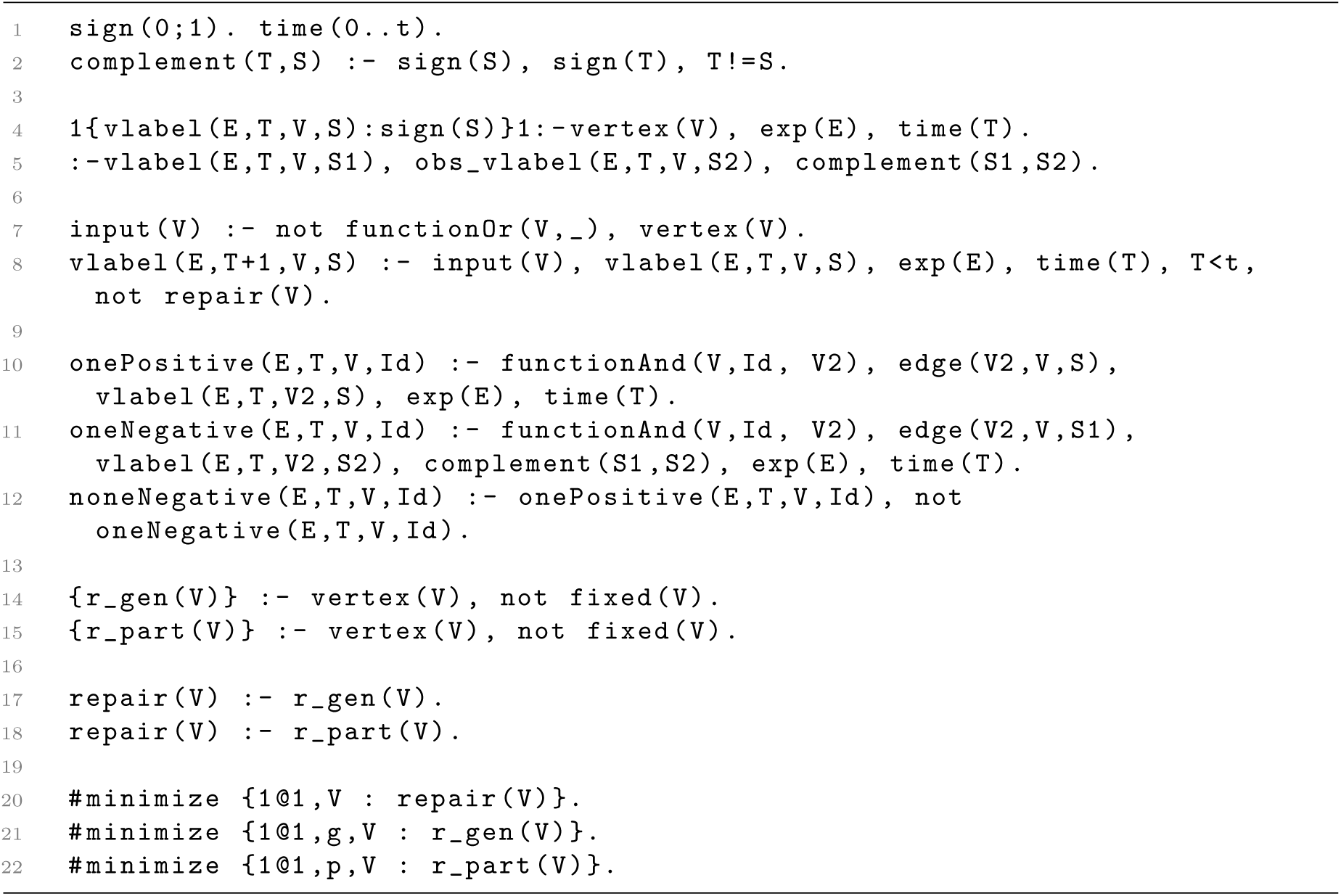
Consistency Check.

A model can have nodes without an associated regulatory function, *i.e.*, nodes without regulators (and no incoming edges in the regulatory graph). We say that these nodes are *input nodes*, as mentioned in Section 2. We define that input nodes maintain the same expected value over time, never changing their value (lines 7 and 8 of Listing 2).

Lines 10, 11, and 12 of Listing 2 define some rules to help compute the result of the regulatory functions. Since regulatory functions are represented as a disjunction of conjunctions, the function is evaluated to true (1) if at least one of the terms of the disjunction is true, and a term is true if all of the literals in the conjunction are true. Therefore, the predicate `onePositive` (resp. `oneNegative`) represents whether there is at least one positive (resp. negative) literal evaluated in a given term. Then, the `noneNegative` predicate represents that none of the literals in a term is negative, meaning that there is a term that evaluates to true, making the output of the regulatory function also true.

Since the application of the regulatory functions depends on the update scheme, the specific update rules are defined in Listing 3 for the synchronous update scheme and in Listing 4 for the asynchronous update scheme.

**Listing 3:**
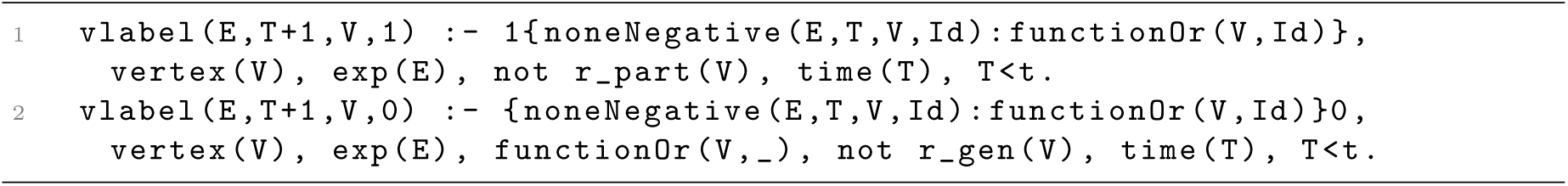
Synchronous Update.

In the synchronous update, all the regulatory functions are applied at each time step, thus updating the corresponding node value. A node is expected to have value 1 in time step `T+ 1` if there is at least one term of its regulatory function that evaluates to true given the expected value of its regulators at time step T (Line 1 of Listing 3). Analogously, a node is expected to have value 0 if none of the terms of its regulatory function evaluates to true (Line 2 of Listing 3).

**Listing 4:**
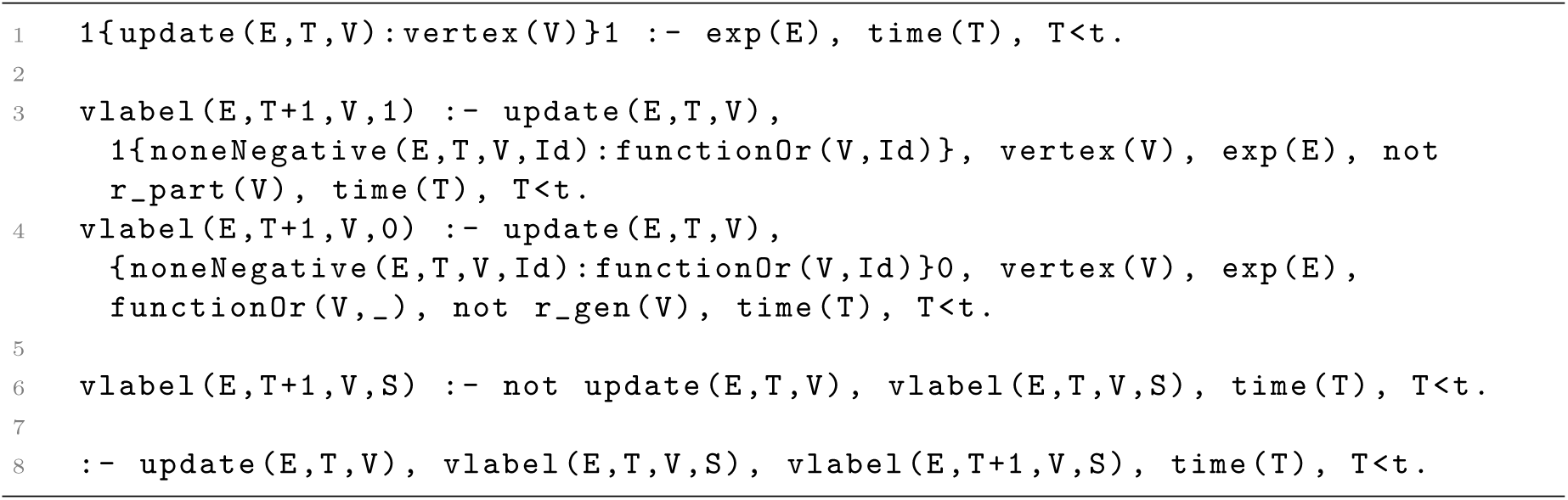
Asynchronous Update.

In the asynchronous update scheme, only one regulatory function is applied at each time step. The predicate `update(E,T,V)` indicates that node `V` is updated at time step `T` in experiment `E`. Line 1 of Listing 4 ensures that at each time step there is one and only one node updated.

The evaluation of the regulatory function (lines 3 and 4 of Listing 4) is similar to the synchronous update scheme rules. However, these rules only apply if the node is updated at the given time step. Therefore, the rule in line 6 defines that if a node is not updated at a given time step then its value is maintained.

Moreover, we add a restriction preventing updates that do not change the state of the network (line 8 of Listing 4).

With the rules described above, it is possible to check the consistency of a model given a set of experimental observations. If a valid *assignment* can be made to all the nodes at every time step, then the model is consistent. Otherwise the model is inconsistent. An *assignment* is valid if the assigned values are equal to the observed values (if they exist) and these values are produced by the correspondent regulatory functions, considering the update scheme in use.

In order to be able to repair an inconsistent model, we add some rules to retrieve information about the reasons of inconsistency. In lines 14-18 of Listing 2 we say that a node can be or not be an inconsistent node in need of repair. Then, in the update rules, we add the information that a node has value 0 (1) if its regulatory function evaluates to false (true) or if it is an inconsistent node in need of repair. This allows a regulatory function not to produce the expected value, and therefore to be considered inconsistent. With this addition, we are able to determine the possible inconsistent nodes in case of an inconsistent model. An optimisation directive is added in lines 20-22 of Listing 2, requiring the minimum number of inconsistent nodes. Moreover, we add some information regarding the type of inconsistency that will be useful when searching for repair operations. We define predicate r_gen(V) meaning that node V is in need of a repair and the expected value is 1 but the function evaluated to 0. The analogous predicate is also defined (r_part(V)) where the expected value is 0 but the function evaluates to 1.

Ultimately, this consistency check procedure analyses a model and a set of experimental observations in time-series data, and determines whether the model is consistent. In case the model is inconsistent, all the possible complete time-series data are considered when missing values in the experimental observation exist. Moreover, this process identifies possible reasons for inconsistency, and produces all possible sets of (minimum) inconsistent nodes and corresponding complete time-series data (as shown in Figure 2).

### 4.2 Model Repair

In order to repair an inconsistent model, four causes of inconsistency are considered, along with their corresponding atomic repair operations, as shown in Table 1.

**Table 1:** Causes of inconsistency and corresponding repair operations

| Cause | Repair Operation |
| --- | --- |
| Wrong Regulatory Function | Function change |
| Wrong Interaction Type | Edge sign flip |
| Wrong Regulator | Edge removal |
| Missing Regulator | Edge addition |

We may have different possible combinations of repair operations that render an inconsistent model consistent. We define a lexicographic optimisation criterion when searching for possible sets of repair operations. This criterion takes into account how a logical model is built. In particular, multiple Boolean functions are likely to be consistent with the existing data, but only one is chosen as the regulatory function of a node. Therefore, we assume that we have a higher level of confidence in the correctness of the topology of the network than in the regulatory functions of the model. Thus, the following lexicographic optimisation criterion is defined with the following order:

1. Minimise the number of add/remove edge operations;
2. Minimise the number of flip sign of an edge operations;
3. Minimise the number of function change operations;

With this optimisation criterion we avoid changing the topology of the network, trying first to apply only function change operations. Then, and only if necessary, trying to flip the sign of an edge. And afterwards, only if still no solution is found, try to add or remove edges to the network.

Given the consistency check procedure previously described in Section 4.1, in case of an inconsistent model it is provided not only all the minimum set of inconsistent nodes, but also the correspondent complete time-series data with the expected values in order to produce a consistent model. With this in mind, we can try to repair each node individually without having to take into account the interactions with other nodes in the network dynamics, as we only have to be sure that each node can produce the expected value at each time step. In order to repair an inconsistent model we consider each and every set of inconsistent nodes and time-series data produced by the consistency check procedure. For each complete expected time-series data and set of inconsistent nodes, we apply the procedure described in Algorithm 1 to each inconsistent node.

#### Algorithm 1 RepairNode(Node)

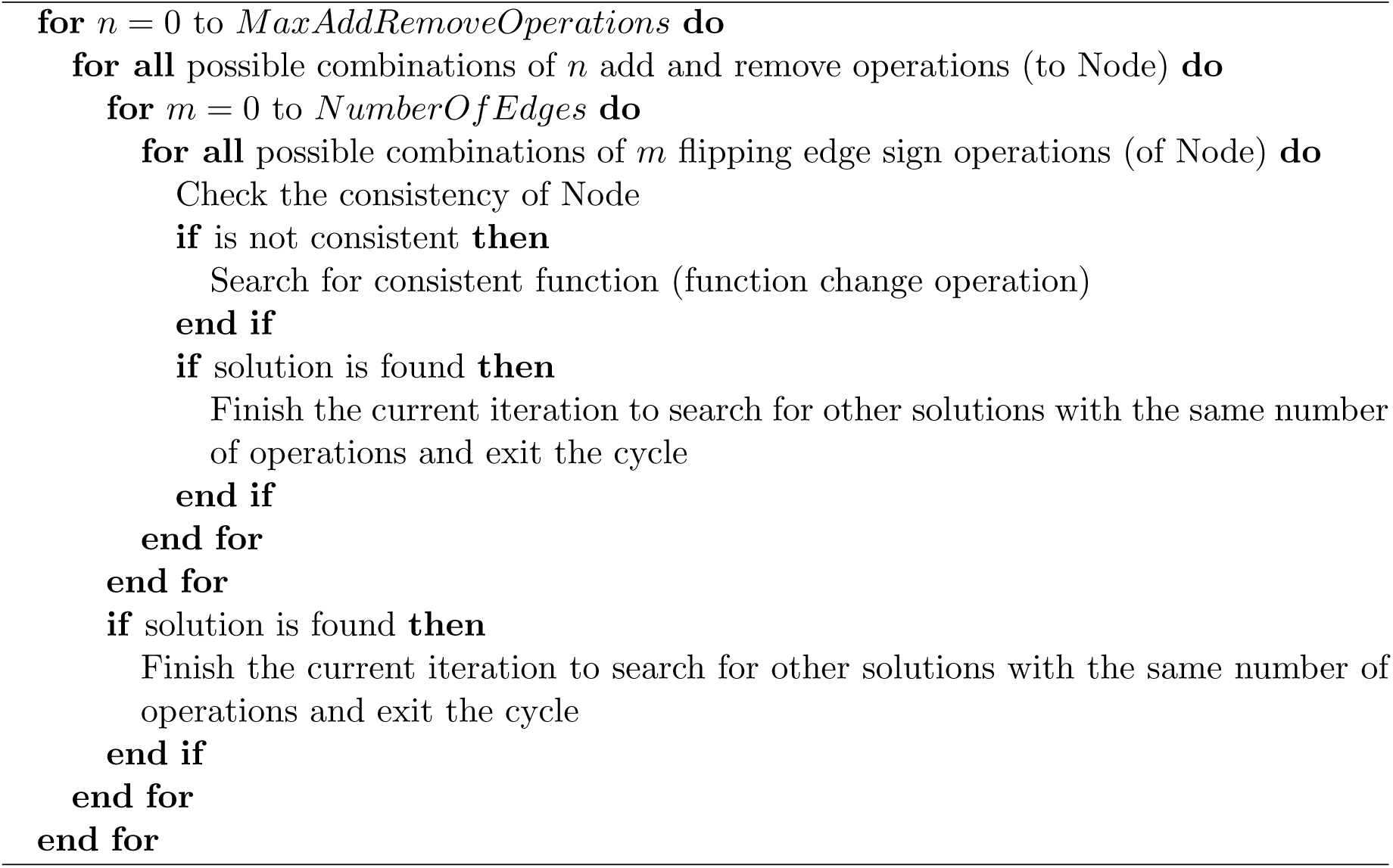

With this procedure, we search for an optimal set of repair operations for each inconsistent node. When repairing a given node we only consider repair operations affecting that node, *i.e.*, we only consider removing edges to that node (removing wrong regulators), adding edges to that node (adding a missing regulator), changing the sign of an edge to that node (changing the type of interaction), and/or changing its regulatory function.

The process of repairing a regulatory function takes advantage of some properties of Boolean functions. In particular, it is possible to define a partial order set over the set of all monotone Boolean functions which can be represented by an Hasse diagram (Crama and Hammer, 2011; Cury *et al.*, 2019). Considering that changing a regulatory function has an impact on the dynamics of the network, it is desirable to have the minimal possible impact. We consider that switching between two closely related functions, with respect to the partial order set, achieves this minimal possible impact (Cury *et al.*, 2019). When repairing an inconsistent function, we therefore search for the set of consistent functions that are closer to the original function, considering the definition of parents and children of monotone Boolean functions as presented by Cury *et al.* (2019).

The previous approach to model revision (Gouveia *et al.*, 2019a,b) had a limitation in the search of possible function repairs. It uses the information regarding the type of inconsistency, as described in Section 4.1, to search for function repairs. If a node has an inconsistency r_part (function produces a 1 but a 0 is expected), a search for more specific functions in the Hasse diagram is performed, starting in the inconsistent function. The process is analogous for the r_gen case. However, if a consistent function is not found, the search for topological repairs starts, considering first flipping the sign of edges operations. This search process does not guarantee that if a consistent function is not found then it does not exist. In fact, a consistent function may exist but may not be an ancestor (or descendant) of the original function, and therefore not found. This leads to a non optimum solution, where a possible function repair could be applied but it is not found and topological repairs are applied instead.

To address this limitation, in this work we implemented an option to force the search of all possible functions when the previous method fails, therefore guaranteeing optimality. However, previous results showed that the search for function repairs has a huge impact on the model revision process as the number of possible functions follows a double exponential with the increase of the number of regulators. Moreover, results show that in most cases consistent functions are actually found with the previous search method. For these reasons, although we give the option to the user to force an exhaustive search, by default we only do this partial search as described in Gouveia *et al.* (2019b). Only when in the presence of inconsistent nodes with both type of inconsistencies, is a more exhaustive search automatically performed.

When repairing a model, with each operation applied, it is necessary to check if the node becomes consistent. To do so, we only need to verify if the expected output is produced by the expected input, taking into account the regulators, interactions and regulatory function. We have to also consider the update scheme used. In a synchronous update, for every time step, it is necessary to verify if considering the current expected values of the regulators, the expected value of the repaired node in the next time step is produced. The node is then considered repaired if it is consistent for every time step. The process is similar for the asynchronous update scheme, for which only the time steps where the repaired node was updated are verified.

This approach produces all the optimal solutions, considering the optimisation criteria defined, in order to render an inconsistent model consistent. However, one may want to prevent some of the repair operations. For example, if one is certain that a given interaction in a model is correct, solutions where that interaction is removed or its type is changed, are not desired. To give the user some control over the solutions produced, it is possible to classify any edge (or even any node) of the network as fixed. With this, solutions that affect fixed edges are not considered, and solutions with fixed nodes marked as inconsistent are not considered. Note that with the addition of such definitions, an inconsistent model may become impossible to repair.

## 5 Experimental Evaluation

Five well-known biological logical models, representative of different processes and organisms, are considered in order to evaluate the proposed tool:

1. The cell-cycle regulatory network of fission yeast (FY) by Davidich and Bornholdt (2008);
2. The segment polarity (SP) network which plays a role in the fly embryo segmentation by Sánchez *et al.* (2002);
3. The T-Cell Receptor (TCR) signaling network by Klamt *et al.* (2006);
4. The core network controlling the mammalian cell cycle (MCC) by Fauré *et al.* (2006);
5. The regulatory network controlling T-helper (Th) cell differentiation by Mendoza and Xenarios (2006).

Table 2 shows some characteristics of the five models used. We use models with 10 to 40 nodes, and different degrees of connectivity. For example, TCR with 40 nodes and 57 edges is less connected than SP, which has the same number of edges but only 19 nodes. More connected networks have, on average, regulatory functions with more regulators.

**Table 2:** Boolean logical models considered for evaluation with corresponding: used abbreviation (Abbr.), number of nodes (#N), number of edges (#E), average number of regulators per node (Avg.Reg.), and maximum number of regulators (Max.Reg.).

| Abbr. | Model | #N | #E | Avg.Reg. | Max.Reg. |
| --- | --- | --- | --- | --- | --- |
| FY | Fission Yeast | 10 | 27 | 3 | 5 |
| SP | Segment Polarity (1 Cell) | 19 | 57 | 3 | 8 |
| TCR | TCR Signalisation | 40 | 57 | 1,425 | 5 |
| MCC | Mammalian Cell Cycle | 10 | 35 | 3,5 | 6 |
| Th | Th Cell Differentiation | 23 | 35 | 1,842 | 5 |

Furthermore, in order to evaluate our tool, we generate corrupted models from the original ones. We consider different types of corruptions: changing a regulatory function (F), flipping a sign of an edge (E), removing a regulator (R), or adding a regulator (A). Considering these corruption types (F, E, R, and A), we generate corrupted models giving a probability of applying each of these operations. We consider 24 different configurations of these parameters. For each configuration, 100 corrupted models are generated, giving a total of 2 400 corrupted models of each original model.

Different sets of observations are also generated. Considering the original models, 10 different observations with 20 time steps for each model are generated. Of these 10 observations, 5 consider the synchronous update scheme and 5 the asynchronous update scheme. The observations generated are consistent with the original models. This allows us to test the proposed approach, by feeding the tool with a corrupted model and the correspondent set of observations, in order to verify if the tool can repair the corrupted model rendering it consistent with the observations given. When testing, we consider different subsets of these generated observations. For each corrupted model we test our tool with only one observation, and with all the five observations simultaneously. Moreover, we consider different lengths of time steps, by considering only observations with three time steps, and with twenty time steps.

For this evaluation we consider the default search of Boolean functions, when searching for function repairs, described in Section 4.2. This may result in non optimum solutions, were the optimal function is not found and topological repairs are applied instead. However, the tool still produces a near optimal solution, considering the optimisation criterion defined. This consideration takes into account that previous results clearly show that the search of function repairs has a big impact on the tool performance. Note that an option to force a complete function search when the default search fails is implemented in the tool, allowing the user to guarantee optimum solutions.

Table 3 shows the median and average times in seconds for solved instances per model, per dynamic update scheme, per number of observations, per number of time steps. Note that for each table entry are considered 2 400 corrupted models. The percentage of solved instances are also shown. All experiments were run on an Intel(R) Xeon(R) 2.1GHz 32-core Linux machine with a time limit of 3 600 seconds (one hour), and a memory limit of 2GB.

**Table 3:**
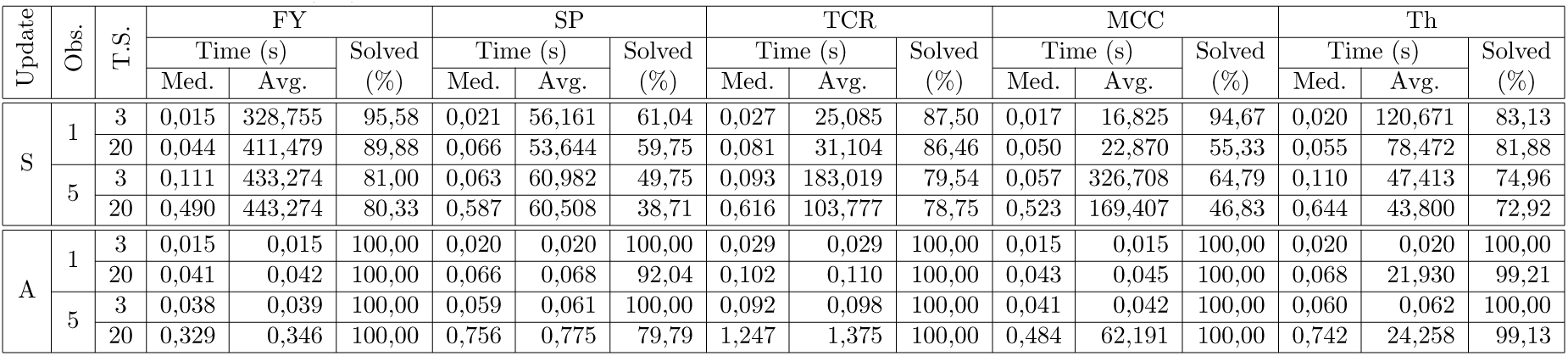
Experimental results. Median (Med.) and average (Avg.) solving times in seconds for each model considering: synchronous (S) and asynchronous (A) update schemes; 1 and 5 observations (Obs.); and 3 and 20 time steps (T.S.) of the dynamic. The percentage of solving instances (%) consider a total of 2400 instances for each table entry.

Table 3 clearly shows that the tool performs better under the asynchronous update scheme than under the synchronous update scheme. This is due to the fact that the synchronous update scheme has more restrictions. Under the synchronous update scheme, all the regulatory functions must produce the expected value of the correspondent node at every time step. Considering that we have one inconsistent node, with five observation with twenty time steps, in the synchronous update scheme, 100 verifications must be performed when repairing that node. However, in the asynchronous update scheme, for each time step only one node is updated, and therefore only that node must be verified. This leads to considerable less verifications when repairing an inconsistent node.

The average solving time increases with the number of experimental observations and with the number of time steps. Conversely, the number of solved instances decreases with the increase of number of observations and time steps. The instances that were not solved were due to time and memory limitations imposed.

Table 3 shows a discrepancy between median and average time of solved instances. Most instances are solved under one second. However, the solving time rapidly grows, as shown in Figure 3.

**Figure 3:**
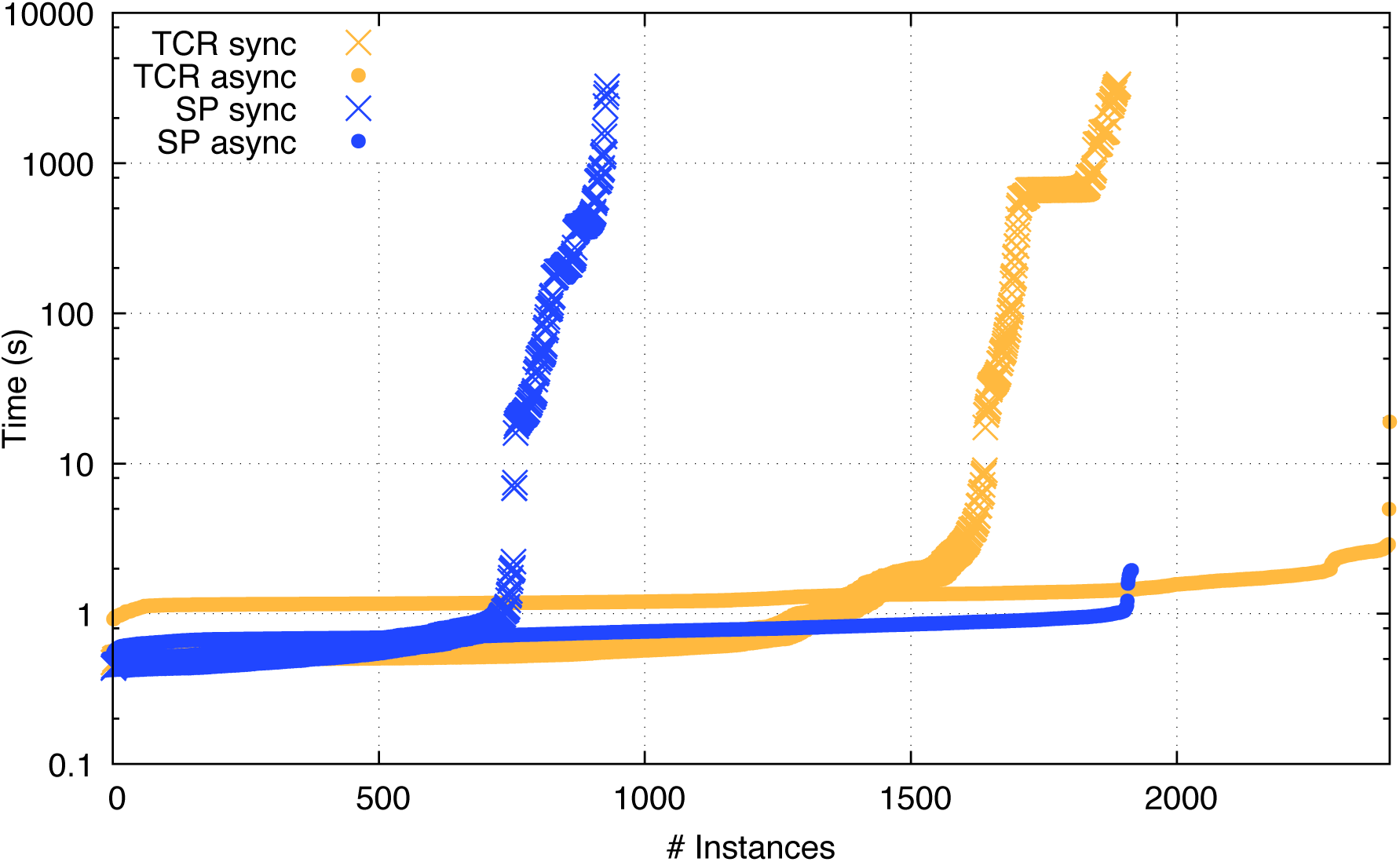
Time comparison of solved instances for the SP and TCR models under synchronous and asynchronous update schemes. Five different observations with twenty time steps were given as input.

The type of corruption made to the original model has a huge impact on the tool performance. Instances where regulators are added or removed take longer to solve. This happens because by adding and/or removing a regulator, the dimension of the correspondent regulatory function is changed. The tool starts by trying to change the regulatory function, which may not be enough, and then considers flipping the sign of the edges. Then, if no solution is found, it considers adding and removing a regulator. Every time the sign of an edge is flipped or a regulator is added or removed, the tool will search for possible function repairs again, compatible with the topological changes. As some operations change the dimension of the function, the search for function repairs must be made from scratch. Moreover, the number of functions grows double exponentially with the number of regulators, which leads to larger search spaces.

Figure 3 shows a comparison between the TCR model (in yellow) and the SP model (in blue). Moreover, it shows both synchronous (crosses) and asynchronous (points) update schemes. The comparison is made considering that five observations are given as input, with twenty time steps each. It is possible to verify that, under the asynchronous update scheme, more instances can be solved. Figure 3 also shows that it is possible to solve more instances of the TCR model than of the SP model. This is due to the fact that SP is a more connected network, having nodes with regulatory functions of greater dimension, leading to bigger search spaces when considering function repairs.

Although we use the default function repair search, not guaranteeing optimum solutions, in the majority of cases, the tool produces solutions with a number of repair operations less than or equal to the number of operations performed to corrupt the model.

## 6 Conclusion

In this work, we propose a model revision tool capable of repairing Boolean logical models with time-series observations under both synchronous and asynchronous dynamics. Our approach verifies the consistency of a given model with a set of observations. In case of inconsistency, all optimal sets of possible repair operations are produced in order to render the model consistent.

The tool was successfully tested using several well-known biological models. In order to test the tool, different time-series data were generated consistent with each model, and different corrupted models were also generated. The tool repairs most corrupted models in less than one second. We conclude that the synchronous update scheme has more restrictions than the asynchronous update leading to greater solving times. This is due to the fact that, in the synchronous case, all the regulatory functions must produce the expected values at every time step.

With this approach we confirm previous results showing that the dimension of the regulatory function has the biggest impact on the tool performance, since the search space of function repairs increases with a double exponential with the number of regulators. For models where the corruption led to repairs considering addition and removal of regulators, the repairing time increased. This phenomenon happens because, when a function dimension changes, not only the tool has to search the correspondent function space, but most probably some topological repair operation occurs, thus changing the function dimension, and performing more function repair searches. This process can have multiple iterations until a consistent repair is found.

Since the function repair search has the biggest impact on the tool performance, we consider a near optimal approach. Although an exhaustive search method is implemented and available to the user, results showed that using the near optimal search approach led to very good results where the number of repair operations produced were less than or equal to the number of operations performed to corrupt the model.

In previous works we addressed the model revision problem under stable state observations (Gouveia *et al.*, 2019a,b). In this work we considered time-series observations under (a)synchronous dynamics, not supporting stable state observations. As future work, we intend to improve this tool in order to support different types of observations simultaneously. It can be of interest, for example, to be able to give a time-series observation indicating that the last state is a stable state.

Furthermore, the proposed tool currently produces all the optimal solutions given the defined optimisation criteria. Heuristics to filter the produced output could also be considered, in particular those proposed in the work of Merhej *et al.* (2017) where rules of thumb are discussed.

## Footnotes

1 Tool available at https://filipegouveia.github.io/ModelRevisionASP

## References

Abou-Jaoudé, W., Traynard, P., Monteiro, P. T., Saez-Rodriguez, J., Helikar, T., Thieffry, D., and Chaouiya, C. (2016). Logical modeling and dynamical analysis of cellular networks. Frontiers in Genetics, 7.

Alchourrón, C. E., Gärdenfors, P., and Makinson, D. (1985). On the logic of theory change: Partial meet contraction and revision functions. J. Symb. Log., 50(2), 510–530.

Biere, A., Heule, M., and van Maaren, H. (2009). Handbook of satisfiability, volume 185. IOS press.

Butte, A. J. and Kohane, I. S. (1999). Mutual information relevance networks: functional genomic clustering using pairwise entropy measurements. In Biocomputing 2000, pages 418–429. World Scientific.

Crama, Y. and Hammer, P. L. (2011). Boolean functions: Theory, algorithms, and applications. Cambridge University Press.

Cury, J. E., Monteiro, P. T., and Chaouiya, C. (2019). Partial Order on the set of Boolean Regulatory Functions. arXiv preprint 1901.07623.

Davidich, M. I. and Bornholdt, S. (2008). Boolean network model predicts cell cycle sequence of fission yeast. PLoS ONE, 3(2), e1672.

De Jong, H. (2002). Modeling and simulation of genetic regulatory systems: a literature review. Journal of computational biology, 9(1), 67–103.

Fauré, A., Naldi, A., Chaouiya, C., and Thieffry, D. (2006). Dynamical analysis of a generic boolean model for the control of the mammalian cell cycle. Bioinformatics, 22(14), e124–e131.

Fayruzov, T., Janssen, J., Vermeir, D., Cornelis, C., and De Cock, M. (2011). Modelling gene and protein regulatory networks with answer set programming. International journal of data mining and bioinformatics, 5(2), 209–229.

Friedman, N. (2004). Inferring cellular networks using probabilistic graphical models. Science, 303(5659), 799–805.

Gebser, M., Guziolowski, C., Ivanchev, M., Schaub, T., Siegel, A., Thiele, S., and Veber, P. (2010). Repair and prediction (under inconsistency) in large biological networks with answer set programming. In KR.

Gouveia, F., Lynce, I., and Monteiro, P. T. (2018). Model revision of logical regulatory networks using logic-based tools. In A. D. Palu’, P. Tarau, N. Saeedloei, and P. Fodor, editors, Technical Communications of the 34th International Conference on Logic Programming (ICLP 2018), volume 64 of OpenAccess Series in Informatics (OASIcs), pages 23:1–23:10, Dagstuhl, Germany. Schloss Dagstuhl–Leibniz-Zentrum fuer Informatik.

Gouveia, F., Lynce, I., and Monteiro, P. T. (2019a). Model revision of boolean regulatory networks at stable state. In Z. Cai, P. Skums, and M. Li, editors, International Symposium on Bioinformatics Research and Applications, pages 100–112. Springer International Publishing.

Gouveia, F., Lynce, I., and Monteiro, P. T. (2019b). Revision of boolean models of regulatory networks using stable state observations. Journal of Computational Biology.

Guerra, J. and Lynce, I. (2012). Reasoning over biological networks using maximum satisfiability. In Principles and Practice of Constraint Programming, pages 941–956. Springer.

Guziolowski, C., Videla, S., Eduati, F., Thiele, S., Cokelaer, T., Siegel, A., and Saez-Rodriguez, J. (2013). Exhaustively characterizing feasible logic models of a signaling network using answer set programming. Bioinformatics, page btt393.

Hopfensitz, M., Müssel, C., Maucher, M., and Kestler, H. A. (2012). Attractors in Boolean networks: a tutorial. Computational Statistics, 28(1), 19–36.

Karlebach, G. and Shamir, R. (2008). Modelling and analysis of gene regulatory networks. Nature Reviews Molecular Cell Biology, 9(10), 770.

Klamt, S., Saez-Rodriguez, J., Lindquist, J. A., Simeoni, L., and Gilles, E. D. (2006). A methodology for the structural and functional analysis of signaling and regulatory networks. BMC Bioinformatics, 7(1), 56.

Lemos, A., Lynce, I., and Monteiro, P. T. (2019). Repairing Boolean logical models from time-series data using Answer Set Programming. Algorithms for Molecular Biology, 14(1), 9.

Mendoza, L. and Xenarios, I. (2006). A method for the generation of standardized qualitative dynamical systems of regulatory networks. Theoretical Biology and Medical Modelling, 3(1), 13.

Merhej, E., Schockaert, S., and De Cock, M. (2017). Repairing inconsistent answer set programs using rules of thumb: A gene regulatory networks case study. International Journal of Approximate Reasoning, 83, 243–264.

Mobilia, N., Rocca, A., Chorlton, S., Fanchon, E., and Trilling, L. (2015). Logical modeling and analysis of regulatory genetic networks in a non monotonic framework. In International Conference on Bioinformatics and Biomedical Engineering, pages 599–612. Springer.

Monteiro, P. T., Abou-Jaoudé, W., Thieffry, D., and Chaouiya, C. (2014). Model checking logical regulatory networks. IFAC Proceedings Volumes, 47(2), 170–175.

Naldi, A., Remy, E., Thieffry, D., and Chaouiya, C. (2011). Dynamically consistent reduction of logical regulatory graphs. Theor. Comput. Sci., 412(21), 2207–2218.

Naldi, A., Monteiro, P. T., Mussel, C., Kestler, H. A., Thieffry, D., Xenarios, I., Saez-Rodriguez, J., Helikar, T., and and, C. C. (2015). Cooperative development of logical modelling standards and tools with CoLoMoTo. Bioinformatics, 31(7), 1154–1159.

Ostrowski, M., Paulevé, L., Schaub, T., Siegel, A., and Guziolowski, C. (2016). Boolean network identification from perturbation time series data combining dynamics abstraction and logic programming. Biosystems, 149, 139–153.

Pauleve, L. (2018). Reduction of qualitative models of biological networks for transient dynamics analysis. IEEE/ACM Transactions on Computational Biology and Bioinformatics, 15(4), 1167–1179.

Sánchez, L., Chaouiya, C., and Thieffry, D. (2002). Segmenting the fly embryo: logical analysis of the role of the segment polarity cross-regulatory module. International Journal of Developmental Biology, 52(8), 1059–1075.

Selvaggio, G., Canato, S., Pawar, A., Monteiro, P. T., Guerreiro, P. S., Brás, M. M., Janody, F., and Chaouiya, C. (2020). Hybrid epithelial-mesenchymal phenotypes are controlled by microenvironmental factors. Cancer Research, page canres.3147.2019.

Siegel, A., Radulescu, O., Le Borgne, M., Veber, P., Ouy, J., and Lagarrigue, S. (2006). Qualitative analysis of the relation between dna microarray data and behavioral models of regulation networks. Biosystems, 84(2), 153–174.

Thomas, R. (1973). Boolean formalization of genetic control circuits. Journal of Theoretical Biology, 42(3), 563–585.

Wegner, I. (1985). The critical complexity of all (monotone) Boolean functions and monotone graph properties. Information and Control, 67(1-3), 212–222.

